# Implementation of a Computing Pipeline for Evaluating the Extensibility of Boolean Networks’ Structure and Function

**DOI:** 10.1101/2020.10.02.323949

**Authors:** Rémi Segretain, Sergiu Ivanov, Laurent Trilling, Nicolas Glade

## Abstract

Formal interaction networks are well suited for representing complex biological systems and have been used to model signalling pathways, gene regulatory networks, interaction within ecosystems, etc. In this paper, we introduce Sign Boolean Networks (SBNs), which are a uniform variant of Threshold Boolean Networks (TBFs). We continue the study of the complexity of SBNs and build a new framework for evaluating their ability to extend, *i.e*. the potential to gain new functions by addition of nodes, while also maintaining the original functions. We describe our software implementation of this framework and show some first results. These results seem to confirm the conjecture that networks of moderate complexity are the most able to grow, because they are not too simple, but also not too constrained, like the highly complex ones. Biological Regulation, Biological Networks, Sign Boolean Networks, Complexity, Extensibility, Network Growth

## 1 Note about this document

This document comes with our article *A Methodology for Evaluating the Extensibility of Boolean Networks’ Structure and Function* published in the *Proceedings of the Complex Networks 2020 conference, Madrid, Spain*[1]. It contains two parts: (i) the description of the manner we implement the computation of our extension problem in a Java-driven pipeline with Answer Set Programming (ASP) calls, (ii) and to the ASP code of the modules used to infer networks.

## 2 Implementation of the extension problem

In section 2.2 of our article [1] we formulate our network extension problem in a logical way, *i.e*. we need to fix a given binary sequence *S_1_* played by at least one node of a SBN N1, a suffix S_k_ (S_1_,S_k_ ∈ {0,1}*), the dimension *d,* and a set of constraints over the quadruplet *(N_1_,S_1_,N,S),* notably the fact that N_1_ is a sub-network of N, and S is a binary sequence played by at least the same node of the sub-network N_1_ plunged into N and is formed by the concatenation of S_1_ and S_k_. As mentioned in the article, such formulation is not effectively usable in its raw form. In practice, we first enumerate exhaustively all networks N1, all their behaviours S1 and all the extended behaviours S, then find all corresponding extended networks N that satisfy the constraints enumerated in section 2.2 of article [1].

In the following, we first give an overview of the software architecture and then show how to solve the crucial issue of inferring efficiently only different networks N.

### 2.1 Overall software architecture

We used a combination of two programming languages, Answer Set Programming (ASP) [3], a non-monotonic logical based programming language, and Java, a classical imperative language. The main software, written in Java, orchestrates the execution and uses multiple ASP modules when needed.

#### 2.1.1 Inference modules in ASP

We first give here a very short introduction to ASP. This logical language allows to express facts and rules, like Prolog, with the help of logical literals. For example, the following rules `p(1). p(2)`. and `c :- p(1), p(2)`. mean that the two facts `p(1)` and `p(2)` are true and that their conjunction implies `c`.

An ASP program infers all logical models (sets of literals) that comply with the facts and rules it specifies: they are called Answer Sets (AS). With the help of *integrity constraints,* logically expressed as rules producing false, some AS can be eliminated. For example, let us consider the two rules `a :- not b. and b :- not a`. ; they accept the two different AS {`a`} and {`b`} (i.e. as `b` cannot be inferred, `b` is considered to be false in ASP, then `not b` is true and `a` is also true). If we add the fact `c`. and the integrity constraint `:- c, not b`. then only the AS {`b, c`} is valid and not {`a, c`}: the integrity constraint discriminates the ASs where the conjunction of `c` and `not b` is be false, then `b` should be true (and `a` is rejected to be true).

The ASP solver that we use, namely *clingcon* [4], proceeds in two steps. First, a *grounder* translates the rules to a propositional form (with only Boolean variables). Then a SAT-like algorithm is applied to this program. We greatly benefit from a recent improvement based on [5] which uses a lazy approach for grounding and then allows the use of numerical variables with a very large range.

In order to increase the flexibility and the re-usability of our ASP code, we cut it into several inference modules dedicated to specific tasks and compatible between each others. Each one introduces a main literal that can be set to configure its module :

- `sbn(N, D)` : implies literals describing a SBN `N` of dimension `D`.
- composedSbn(N_*a*_, N_*b*_) : constrains two given SBNs N_*a*_ and N_*b*_ so that one SBN is an extension of the other as defined in section 2.2 of [1].
- `sequencePlayedBySbn(N, S, I)` : constrains a SBN `N` to play a given sequence `S` on his node of index `I`.
- `orderedSbn(N)` : from a given SBN `N`, generates the equivalent ordered one by permutation of nodes (see section 2.2.3).

#### 2.1.2 Processing Pipeline

The main software is organized as a pipeline of processors (called jobs): each job, programmed in Java, does its own part of the work, then furnishes its results to the next job, *etc*. Using a custom Java library [2], the workload is divided into tasks that we scatter across the different jobs, allowing to make the execution parallel on any number of CPU cores, thus enhancing the overall performances. An overview of this pipeline is given figure 1. The potential of the combination of an imperative language and inference modules lay in the facts that we can take advantages from both: efficiently find data sets by ASP inference, filter and enrich them in the Java pipeline. Moreover, those enriched data set can be used in return to configure other calls to inference modules. This could not have been easily done using only ASP.

**Figure 1:**
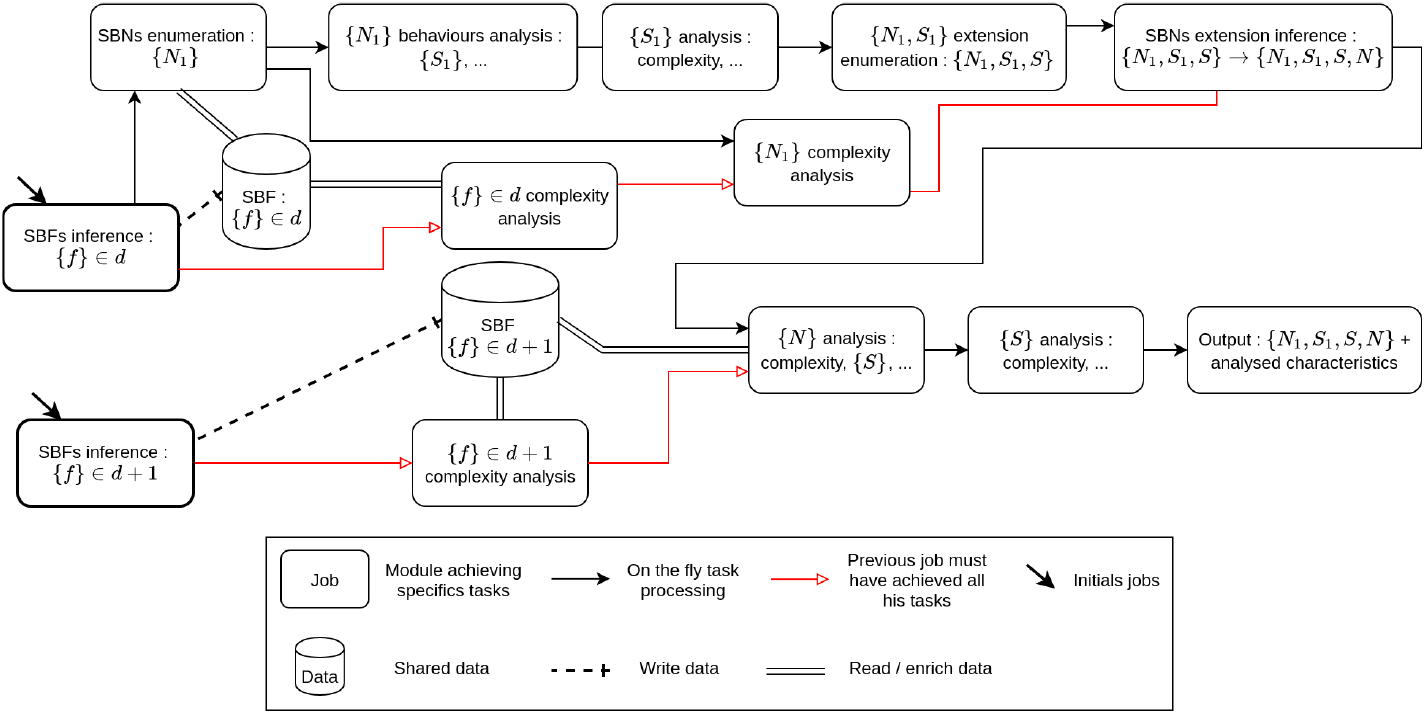
Overview of the processing pipeline, jobs division and ordering. Initial jobs use inference modules to find the SBFs, which are used to generate the SBNs. Then, all SBN behaviors are analysed. The complexity of individual SBFs, SBNs and behaviors is determined and every reachable extended behaviors are listed (the program generates sequence with *S*_1_ as sub-sequence). At this point we have got all the triplets (*N*_1_, *S*_1_, *S*). For each of them, inference modules are called another time to find all the extended networks *N*.

The jobs use inference modules by generating literals that link the modules they need. In practice, such a job generates an ASP file that imports and configures the needed modules. Here is a casual example: a job knows a SBF *f* and aims to find all SBNs that contain *f* and play the sequence *s* = (100)*. This job configures the necessary inference modules this way:

~~~
//implies the generation of a SBN n of dimension 3 sbn (n, 3).
// constrains n to contain f
:– not sbf (n, f).
// constrain n to play the sequence s
:– not sequencePlayedBySbn (n, s).
// define the bits of s and their order
sequence(s, 1, 1). // the first element of s is 1.
sequence (s, 2, 0).
sequence (s, 3, 0).
~~~

To infer ASs different only on specified predicates, jobs that call inference modules take advantage of an option of the solver: *project*, If several AS are formed by the sames atoms belonging to a list of literals, this option will force the program to keep only one of those AS. For example, let us consider the two following AS AS_1_ : {p(a), q(b), r(c)} and AS_2_ :{ p(a), q(b), r(d)}. The projection on `p(X)` and `q(X)` will only provide us either AS_1_ or AS_2_ as they both contains the same literals for `p(X)` and `q(X)`.

### 2.2 SBF and SBN generation issues

When generating SBF and SBN we encountered several issues that needed particular implementation solutions:

1. How to find all unique SBFs of dimension *d* ? Within multiple sets of weights like {*w*_1_, *w*_2_, *w*_3_} to define several SBFs, their truth table output may be equal or equivalent through inputs permutation, so they belong to the same equivalence class 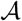. To avoid duplicates, we must use only one representative per equivalence class. The core problem is then *how can we found all those equivalence classes reliably ?*
2. How to enumerate only unique SBNs from those SBFs ?

a. Given a set of *d* SBFs, redundant SBNs can be enumerated by way of permutation of the SBFs over the nodes, *e.g. f*_1_ associated to node 1 and *f*_2_ associated to node 2 or the alternative.
b. There are also two others sources of variations that produce the same sets of SBNs. Both correspond to layout variations, but they are obtained in two different ways:

- Permutation of the source node for the inputs of the SBF, *e.g*. input 1 from node 1 and input 2 from node 2 or the alternative.
- Two SBFs can also be obtained from each other through permutation of their input weights. For example *Weights*(*f*_1_) = 〈−1, 2, 1〉 is equivalent to *Weights*(*f*_2_) = 〈2, −1, 1〉 by permutation of *w*_1_ and *w*_2_. As we must avoid the exploration of duplicate SBNs, which would alter the results and be very costly in computational power, we must take into account only one of these variations in our SBN enumeration. For performance issues detailed below, we choose to keep the layout variation due to input’s source node permutations and neutralize the other.

#### 2.2.1 Inference of SBFs

Satisfiable answer sets of SBFs are generated using both an inference module and job post-processing. During these operations, we must generate only one SBF per equivalence class and neutralize the redundancy induced by input weight permutations.

Within the inference module in ASP, a SBF is addressed by its abstract form 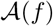. Thanks to the *project* option of *clingcon* (see remark 2.1.2 above) the SBFs from different layouts 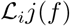 but belonging to the same 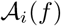 are regrouped in only one equivalent class 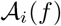. Since a 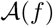 is by definition a set of constraints over the SBF weights, it is particularly easy to specify 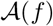 in ASP, particularly when using *integer linear constraints*, a new ASP improvement. We can then infer a set of SBFs classified by 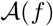 and by the sum of absolutes values of weights. Once inference of SBFs is done, Java keeps only one minimal representative in each equivalence class (see article [1], section 2.1).

The generated set of SBFs solve the two issues 1 and 2b presented in 2.2. By dealing with the 2b issue from the beginning, we limit the number of processed SBFs in the future jobs, saving both CPU time and RAM consumption. In addition of the minimal representative, several others SBFs may be kept from an equivalent class if they have weights set to zero. Edges with *w* = 0 are considered as non-existing edges. In consequence, even if two SBFs belong to the same equivalence class, different directed graphs can be obtained when some edges are absent. As the structural complexity is based on the topology of the interaction graph, this lead to different and unique SBNs that we must also explore.

#### 2.2.2 Order over SBF

Using 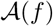 makes it possible to determine an order over the SBFs.

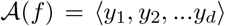 is composed of a unique ordered set of numerical values (*y_i_*), each value bounded between 0 and the number of configurations *Card*(*X_i_*) of *X* in *Nai_i_*. Each digit of the vector 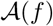 is encoded using a different numerical base *Base_i_* in such a way this vector could be converted into a number in decimal base 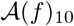. The numerical base corresponds to the number of configurations *Card*(*X_i_*) of *X* in *Nai_i_*, increased by 1 to include 0 (*y_i_* ∈ [0; *Card*(*X_i_*)]) : *Base_i_* = *Card*(*X_i_*) + 1.

In 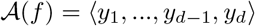, *y_d_* code for the units, *y*_*d*−1_ for the decades, *y*_*d*−2_ for the hundreds, *etc*, so it matches with the conventional order in which number are read. To convert 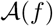 into a 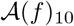 we have:

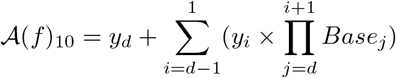

Every different 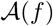 of the same dimension will be converted into a unique 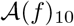, as illustrated with 2 SBFs in table 1, that we can use to define a order over SBFs such that :

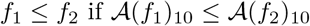

**Table 1:** Examples of conversion of 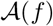 into a 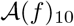 with 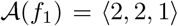 and 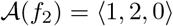

| | $y_1$ | $y_2$ | $y_3$ | $\mathcal{A}(f_1)_{10}$ | | $y_1$ | $y_2$ | $y_3$ | $\mathcal{A}(f_2)_{10}$ |
| --- | --- | --- | --- | --- | --- | --- | --- | --- | --- |
| $\mathcal{A}(f_1)$ | 2 | 2 | 1 | 21 | $\mathcal{A}(f_2)$ | 1 | 2 | 0 | 12 |
| $Card(X_i)$ | 3 | 3 | 1 | | $Card(X_i)$ | 3 | 3 | 1 | |
| $Base_i$ | 4 | 4 | 2 | | $Base_i$ | 4 | 4 | 2 | |

#### 2.2.3 SBN enumeration

In order to avoid the generation of multiple SBN composed of the same SBFs but assigned to different nodes (issue 2a in section 2.2.3), we use the order over SBFs and follow a simple rule to assign the SBFs to labelled nodes, the one after the other: the unassigned SBF with the lower 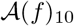 is always linked to the unassigned node with the lower index, and so forth.

The last step is the construction of the network layout, *i.e*. the assignation of a source node to each SBF input. For a given dimension, the number of different layouts is fixed and equal to *d*!^*d*^. Figure 2 gives the 4 different layouts available in dimension 2 and shows how nodes are assigned to SBF inputs.

**Figure 2:**
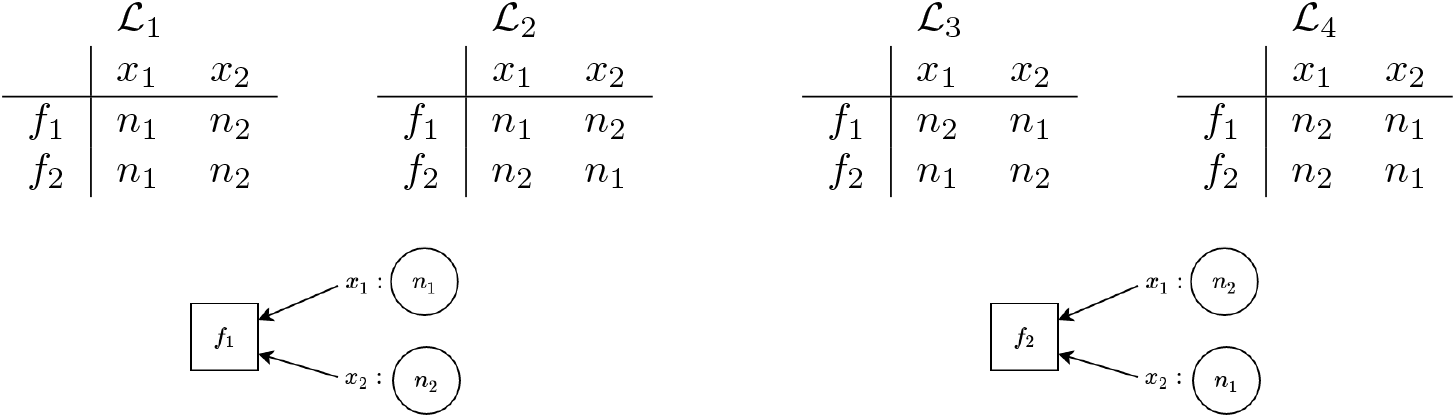
List of the 4 possible layouts for 2-dimension SBNs: 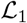 to 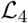. There is a directed edge going from the node *n_i_* to the node associated to the SBF *f_i_* when an input *x_i_* of *f_i_* reads the value of *n_i_*. The two diagrams below illustrate the input assignments of SBFs *f*_1_ and *f*_2_ to nodes *n*_1_ and *n*_2_ according to layout 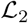.

The generation of layouts is independent to both the assignment of the SBFs *f_i_* to the nodes *n_i_*, and that of the nodes *n_i_* to the inputs *x_i_* of SBFs. Consequently we only need to generate these layouts once before SBF-node assignment. This lead to a huge saving in CPU time and RAM consumption.

Once all layouts are obtained, we combine every composition SBFs/nodes to all layouts to finally generate all possible SBNs. In the end we obtain the set of all unique SBN such that they are built with minimal weights and that there is no other SBN figuring the same set of SBFs with the same layout. As zero-weighted edges are considered absent, we may obtain networks figuring disconnected nodes. Those “networks” are not considered as functional networks, so they are discarded.

## 3 Inference modules in ASP

The following sections contain the ASP code of the modules listed in 2.1.1.

### 3.1 SBN

~~~
%∗
 ∗ Constructs all the possibilities for the network sbn(Name, Dimension)
 ∗ At least one predicate of this from must exist.
 ∗%
% means that constraints operated by clingcon 2017 are involved.
#include <csp>.
%% LIMIT CONSTRAINTS
maxDim(4).
possibleDim(1..MaxDim) :- maxDim(MaxDim).
possibleIdx(1..MaxDim) :- maxDim(MaxDim).
:-
    not sbn(_, _)
.
%% NETWORK GENERATION
% Generation of possible indices for a network based on the dimension
possibleNetworkIndex(NetworkName, Idx) :-
    sbn(NetworkName, Dimension)
  , possibleDim(Dimension)
  , Idx > 0
  , Idx <= Dimension
  , possibleIdx(Idx)
.
% Generation of network nodes
node(NetworkName, Idx) :-
    sbn(NetworkName, Dimension)
  , possibleNetworkIndex(NetworkName, Idx)
.
% Generation of node inputs
{
   nodeInput(NetworkName, NodeIdx, InputIdx, SrcNodeIdx) :
       node(NetworkName, NodeIdx)
     , node(NetworkName, SrcNodeIdx)
     , possibleNetworkIndex(NetworkName, InputIdx)
}.
% A node only has a single input coming from any other given node
:-
    nodeInput(NetworkName, NodeIdx, InputIdx1, SrcNodeIdx)
  , nodeInput(NetworkName, NodeIdx, InputIdx2, SrcNodeIdx)
  , InputIdx1 != InputIdx2
.
% A node receives any given input from a single other node
% (an input arc cannot originate at more than one source node)
:-
   nodeInput(NetworkName, NodeIdx, InputIdx, SrcNodeIdx1)
  , nodeInput(NetworkName, NodeIdx, InputIdx, SrcNodeIdx2)
  , SrcNodeIdx1 != SrcNodeIdx2
.
% For every node, an input coming from every other node in the network must exist.
% Therefore, Dimension^2 nodeInputs must exist for each network.
:-
    #count{ NodeIdx, InputIdx, SrcNodeIdx :
       nodeInput(NetworkName, NodeIdx, InputIdx, SrcNodeIdx)
    } != Dimension∗∗2
  , sbn(NetworkName, Dimension)
.
% Generation of weights for every input of every node
&dom{-Dimension..Dimension} = weight(NetworkName, NodeIdx, InputIdx) :-
node(NetworkName, NodeIdx)
, possibleNetworkIndex(NetworkName, InputIdx)
  , sbn(NetworkName, Dimension)
.
% Enumeration of the inequalities of the functions implemented by the nodes
{
ineq(NetworkName, I, Input1) :
     node(NetworkName, I)
, possibleNetworkIndex(NetworkName, Input1)
;ineq(NetworkName, I, Input1, Input2) :
      node(NetworkName, I)
    , possibleNetworkIndex(NetworkName, Input1)
    , possibleNetworkIndex(NetworkName, Input2)
, Input1 < Input2
;ineq(NetworkName, I, Input1, Input2, Input3) :
      node(NetworkName, I)
    , possibleNetworkIndex(NetworkName, Input1)
    , possibleNetworkIndex(NetworkName, Input2)
    , possibleNetworkIndex(NetworkName, Input3)
, Input1 < Input2
, Input2 < Input3
;ineq(NetworkName, I, Input1, Input2, Input3, Input4) :
      node(NetworkName, I)
    , possibleNetworkIndex(NetworkName, Input1)
    , possibleNetworkIndex(NetworkName, Input2)
    , possibleNetworkIndex(NetworkName, Input3)
    , possibleNetworkIndex(NetworkName, Input4)
, Input1 < Input2
, Input2 < Input3
, Input3 < Input4
}.
% Constraints between the weights and the inequalities
&sum{weight(NetworkName, NodeIdx, Input1)} > 0 :-
   ineq(NetworkName, NodeIdx, Input1)
.
&sum{weight(NetworkName, NodeIdx, Input1)} < = 0 :-
     not ineq(NetworkName, NodeIdx, Input1)
   , node(NetworkName, NodeIdx)
   , possibleNetworkIndex(NetworkName, Input1)
.
&sum{weight(NetworkName, NodeIdx, Input1); weight(NetworkName, NodeIdx, Input2)} > 0 :-
    ineq(NetworkName, NodeIdx, Input1, Input2)
.
&sum{weight(NetworkName, NodeIdx, Input1); weight(NetworkName, NodeIdx, Input2)} <= 0 :-
    not ineq(NetworkName, NodeIdx, Input1, Input2)
  , node(NetworkName, NodeIdx)
  , possibleNetworkIndex(NetworkName, Input1)
  , possibleNetworkIndex(NetworkName, Input2)
  , Input1 < Input2
.
&sum{weight(NetworkName, NodeIdx, Input1); weight(NetworkName, NodeIdx, Input2); weight(NetworkName, NodeIdx, Input2); weight(NetworkName, NodeIdx, Input3)} > 0 :-
    ineq(NetworkName, NodeIdx, Input1, Input2, Input3)
.
&sum{weight(NetworkName, NodeIdx, Input1); weight(NetworkName, NodeIdx, Input2); weight(NetworkName, NodeIdx, Input3)} <= 0 :-
    not ineq(NetworkName, NodeIdx, Input1, Input2, Input3)
  , node(NetworkName, NodeIdx)
  , possibleNetworkIndex(NetworkName, Input1)
  , possibleNetworkIndex(NetworkName, Input2)
  , possibleNetworkIndex(NetworkName, Input3)
  , Input1 < Input2
  , Input2 < Input3
.
&sum{weight(NetworkName, NodeIdx, Input1); weight(NetworkName, NodeIdx, Input2); weight(NetworkName, NodeIdx, Input4)} > 0 :-
    ineq(NetworkName, NodeIdx, Input1, Input2, Input3, Input4)
.
&sum{weight(NetworkName, NodeIdx, Input1); weight(NetworkName, NodeIdx, Input2); weight(NetworkName, NodeIdx, Input4)} <= 0 :-
    not ineq(NetworkName, NodeIdx, Input1, Input2, Input3, Input4)
  , node(NetworkName, NodeIdx)
  , possibleNetworkIndex(NetworkName, Input1)
  , possibleNetworkIndex(NetworkName, Input2)
  , possibleNetworkIndex(NetworkName, Input3)
  , possibleNetworkIndex(NetworkName, Input4)
  , Input1 < Input2
  , Input2 < Input3
  , Input3 < Input4
.
% Determines the components of the abstract representations of the functions
% implemented by the nodes.
% sbf(NetworkName, NodeIdx, IneqInputs, IneqCount)
sbf(NetworkName, NodeIdx, 1, X) :-
  #count{
Input1 :
ineq(NetworkName, NodeIdx, Input1)
} = X
  , node(NetworkName, NodeIdx)
  , sbn(NetworkName, Dimension)
  , Dimension >= 1
.
sbf(NetworkName, NodeIdx, 2, X) :-
  #count{
Input1, Input2 :
ineq(NetworkName, NodeIdx, Input1, Input2)
}=X
  , node(NetworkName, NodeIdx)
  , sbn(NetworkName, Dimension)
  , Dimension >= 2
.
sbf(NetworkName, NodeIdx, 3, X) :-
  #count{
Input1, Input2, Input3 :
ineq(NetworkName, NodeIdx, Input1, Input2, Input3)
}=X
  , node(NetworkName, NodeIdx)
  , sbn(NetworkName, Dimension)
  , Dimension >= 3
.
sbf(NetworkName, NodeIdx, 4, X) :-
#count{
Input1, Input2, Input3, Input4 :
ineq(NetworkName, NodeIdx, Input1, Input2, Input3, Input4) }=X
  , node(NetworkName, NodeIdx)
  , sbn(NetworkName, Dimension)
  , Dimension >= 4
.
% There must exist as many SBFs as there are nodes.
    #count{ NodeIdx, IneqInputs :
          sbf(NetworkName, NodeIdx, IneqInputs, _)
    } != Dimension∗∗2
  , sbn(NetworkName, Dimension)
.
% Compute the factorial values needed for the SBF/Node assignation
factorial(0, 1).
factorial(1, 1).
maxFactorial(MaxFactorial) :- maxDim(MaxFactorial).
factorial(X, Value1) :-
    Value1 = X ∗ Value2
  , factorial(X-1, Value2)
  , maxFactorial(Max)
  , X <= Max
.
% Compute the maximum number of configuration in each ineqInputs category (Nai)
maxIneq(NetworkName, IneqInputs, MaxValue) :-
    sbn(NetworkName, Dimension)
  , sbf(NetworkName, _, IneqInputs, _)
  , factorial(Dimension, FactDim)
  , factorial(IneqInputs, FactIneqInputs)
  , factorial(Dimension - IneqInputs, FactDimMinusIneqInputs)
  , MaxValue = (FactDim/(FactIneqInputs ∗ FactDimMinusIneqInputs))
.
% compute the numerical base of each category if IneqInputs (Nai)
weightNbIneq(NetworkName, IneqInputs, Weight) :-
   maxIneq(NetworkName, IneqInputs, MaxValue)
  , sbn(NetworkName, Dimension)
  , IneqInputs = Dimension
  , Weight = (MaxValue+1)
.
weightNbIneq(NetworkName, IneqInputs, Weight) :-
    maxIneq(NetworkName, IneqInputs, MaxValue)
  , weightNbIneq(NetworkName, IneqInputs+1, WeightPrec)
  , Weight = (MaxValue+1) ∗ WeightPrec
.
% compute the SBF decimal “value”
nodeFunctionValue(NetworkName, NodeIdx, FunctionValue) :-
    #sum{ NbIneq ∗ Weight :
        sbf(NetworkName, NodeIdx, IneqInputs, NbIneq)
      , weightNbIneq(NetworkName, IneqInputs, Weight)
    } = FunctionValue
  , node(NetworkName, NodeIdx)
.
% ensure that the attribution of the SBFs over the nodes follow the SBF order and the node index order.
:-
    nodeFunctionValue(NetworkName, NodeIdx1, FunctionValue1)
  , nodeFunctionValue(NetworkName, NodeIdx2, FunctionValue2)
  , NodeIdx1 < NodeIdx2
  , FunctionValue1 > FunctionValue2
  , orderedNodeFunction(NetworkName)
.
~~~

### 3.2 SBN Composition

~~~
%%% Constraint a network to be composed by a other one.
%%% composedSbn(Sbn1, Sbn2) : Sbn1 is composed of Sbn2
%%% Can be use only once per call in this actual form
% means that constraints operated by clingcon 2017 are involved.
#include <csp>.
% force the presence of this predicates to use the module
:-
  not composedSbn(_, _)
.
% there can be only one use of this module per call
:-
    not #count{ Sbn1, Sbn2 : composedSbn(Sbn1,Sbn2) } = 1
.
% the given SBNs must exists
:-
    composedSbn(Sbn1, Sbn2)
  , not sbn(Sbn1, _)
.
:-
    composedSbn(Sbn1, Sbn2)
  , not sbn(Sbn2, _)
.
% The dimensions of the given SBNs must be compatible
:-
    composedSbn(Sbn1, Sbn2)
  , sbn(Sbn1, Dimension1)
  , sbn(Sbn2, Dimension2)
  , not Dimension1 >= Dimension2
.
% creation of a subnetwork called “extrusion” include in Sbn1
sbn(extrusion, Dimension) :-
    composedSbn(Sbn1, Sbn2)
  , sbn(Sbn2, Dimension)
.
% constraint the extrusion weights to be the same as Sbn1
:-
    sbn(extrusion, Dimension)
  , &sum{weight(Sbn1, NodeIdx, InputIdx)} = W1
  , &sum{weight(extrusion, NodeIdx, InputIdx)} = W2
  , not W1 = W2
  , limitWeights(Sbn1, W1)
  , limitWeights(Sbn1, W2)
  , possibleNetworkIndex(extrusion, NodeIdx)
  , possibleNetworkIndex(extrusion, InputIdx)
.
% constraint the structure of the extrusion to be the same as Sbn1 and Sbn2
:-
  sbn(extrusion, Dimension)
  , composedSbn(Sbn1, Sbn2)
  , sbn(Sbn2, Dimension)
  , nodeInput(extrusion, NodeIdx, InputIdx, SrcNodeIdx1)
  , nodeInput(Sbn2, NodeIdx, InputIdx, SrcNodeIdx2)
  , not SrcNodeIdx1 = SrcNodeIdx2
.
% constraint the SBFs of the extrusion to be the same as Sbn2
:-
    sbn(extrusion, Dimension)
  , composedSbn(Sbn1, Sbn2)
  , sbn(Sbn2, Dimension)
  , sbf(extrusion, NodeIdx, NbIneqInputs, IneqCount1)
  , sbf(Sbn2, NodeIdx, NbIneqInputs, IneqCount2)
  , not IneqCount1 = IneqCount2
.
% specify the weight limit for a SBN according to his dimension
limitWeights(NetworkName, -Dimension..Dimension) :-
    sbn(NetworkName, Dimension)
.
~~~

### 3.3 Sequence played by SBN

~~~
%%% Check the network behavior to match a given music (sequence) on a given node.
% means that constraints operated by clingcon 2017 are involved.
#include <csp>.
% forbid the use of the module without the presence of this predicate
:-
    not musicPlayBySbn(_, _, _)
.
% the given network and music must exists
:-
    musicPlayBySbn(_, MusicName, _)
  , not music(MusicName, _, _)
.
:-
    musicPlayBySbn(NetworkName, _, _)
  , not sbn(NetworkName, _)
.
% the given node must exist in the network
:-
    musicPlayBySbn(NetworkName, _, NodeIdx)
  , sbn(NetworkName, Dimension)
  , not possibleNetworkIndex(NetworkName, NodeIdx)
.
% the music length must fit with the network dimension
:-
  musicPlayBySbn(NetworkName, MusicName, _)
  , sbn(NetworkName, Dimension)
  , musicSize(MusicName, MusicSize)
  , not MusicSize <= Dimension∗∗2
.
% tell the music length
musicSize(MusicName, Size) :-
music(MusicName, _, _)
, Size = {music(MusicName, _, _)}
.
% give the possible index for the note of the music
musicStep(MusicName, StepIdx+1) :-
musicSize(MusicName, StepIdx)
.
musicStep(MusicName, StepIdx) :-
StepIdx > 0
, musicStep(MusicName, StepIdx+1)
.
% give the State of node NodeIdx at the step StepIdx of a music MusicName : nodeState(NodeIdx, MusicName, StepIdx, State)
% initialise the first state
1{nodeState(NetworkName, NodeIdx, MusicName, 1, 0);nodeState(NetworkName, NodeIdx, MusicName, MusicName, 1, 1)}1 :-
possibleNetworkIndex(NetworkName, NodeIdx)
, musicStep(MusicName, 1)
.
% transfom the enumerated weight into predicates
limitWeights(NetworkName, -Dimension..Dimension) :-
    sbn(NetworkName, Dimension)
.
inputWeightForStep(NetworkName, NodeIdx, MusicName, InputIdx, StepIdx, Value) :-
State = 1
, &sum{weight(NetworkName, NodeIdx, InputIdx)} = Value
, limitWeights(NetworkName, Value)
, nodeInput(NetworkName, NodeIdx, InputIdx, SrcNodeIdx)
, nodeState(NetworkName, SrcNodeIdx, MusicName, StepIdx-1, State)
.
inputWeightForStep(NetworkName, NodeIdx, MusicName, InputIdx, StepIdx, 0) :-
State = 0
, nodeInput(NetworkName, NodeIdx, InputIdx, SrcNodeIdx)
, nodeState(NetworkName, SrcNodeIdx, MusicName, StepIdx-1, State)
.
% specify the state of a node at a given step
nodeState(NetworkName, NodeIdx, MusicName, StepIdx, 1) :-
  musicStep(MusicName, StepIdx)
, musicStep(MusicName, StepIdx-1)
, #sum{Value, InputIdx : inputWeightForStep(NetworkName, NodeIdx, MusicName, InputIdx, StepIdx, Value)} ~ 0
  , sbn(NetworkName, _)
  , possibleNetworkIndex(NetworkName, NodeIdx)
.
nodeState(NetworkName, NodeIdx, MusicName, StepIdx, 0) :-
  musicStep(MusicName, StepIdx)
, musicStep(MusicName, StepIdx-1)
, #sum{Value, InputIdx : inputWeightForStep(NetworkName, NodeIdx, MusicName, InputIdx, StepIdx, Value)} <= 0
  , sbn(NetworkName, _)
  , possibleNetworkIndex(NetworkName, NodeIdx)
.
% force the state of the playing node to match the music
:-
  nodeState(NetworkName, NodeIdx, MusicName, StepIdx, NodeState)
  , musicPlayBySbn(NetworkName, MusicName, NodeIdx)
  , music(MusicName, StepIdx, MusicNote)
  , musicSize(MusicName, MusicSize)
  , StepIdx
  , NodeState != MusicNote
.
% specify the network state at a given step : networkState(MusicName, StepIdx, State)
possibleState(NetworkName, 0..(2∗∗Dimension)) :-
    sbn(NetworkName, Dimension)
.
networkState(NetworkName, MusicName, StepIdx, State) :-
State = #sum{(2∗∗(NodeIdx-1))∗ NodeState :
nodeState(NetworkName, NodeIdx, MusicName, StepIdx, NodeState)
}
, possibleState(NetworkName, State)
, musicStep(MusicName, StepIdx)
.
% ensure different network state for each step of the music
:-
networkState(NetworkName, MusicName, StepIdx1, State)
  , networkState(NetworkName, MusicName, StepIdx2, State)
  , musicPlayBySbn(NetworkName, MusicName, _)
  , musicSize(MusicName, MusicSize)
  , StepIdx1 <= MusicSize
  , StepIdx2 <= MusicSize
, StepIdx1 != StepIdx2
.
% ensure the path along the transition graph is a cycle
:-
networkState(NetworkName, MusicName, 1, State1)
  , networkState(NetworkName, MusicName, MusicSize+1, State2)
  , musicPlayBySbn(NetworkName, MusicName, _)
  , musicSize(MusicName, MusicSize)
, State1 != State2
.
~~~

### 3.4 Ordered version of SBN

~~~
% means that constraints operated by clingcon 2017 are involved.
#include <csp>.
% forbid the use of the module without this predicate
:-
  not generateOrderedFunctionsVersionOfSbn(_)
.
% the concerned network must exist
:-
    generateOrderedFunctionsVersionOfSbn(NetworkName)
  , not sbn(NetworkName, _)
.
% map the function over the node
% save the index mapping
2{orderedNodeFunctionValue(NetworkName, NewNodeIdx+1, FunctionValue); mapIndex(NetworkName, OldNodeIdx, NewNodeIdx+1)}2 :-
    #count{ NodeIdx, ValueX :
        nodeFunctionValue(NetworkName, NodeIdx, ValueX)
     , ValueX < FunctionValue
     , NodeIdx != OldNodeIdx
   } = NewNodeIdx
, possibleNetworkIndex(NetworkName, NewNodeIdx+1)
, nodeFunctionValue(NetworkName, OldNodeIdx, FunctionValue)
, sbn(NetworkName, _)
, generateOrderedFunctionsVersionOfSbn(NetworkName)
, not nodeFunctionValue(NetworkName, OldNodeIdxB, FunctionValue) :
   OldNodeIdxB < OldNodeIdx
   , possibleNetworkIndex(NetworkName, OldNodeIdxB)
.
2{orderedNodeFunctionValue(NetworkName, NewNodeIdx + Shift + 1, FunctionValue); mapIndex(NetworkName, OldNodeIdx, NewNodeIdx + Shift + 1)}2 :-
    #count{ NodeIdx, ValueX :
        nodeFunctionValue(NetworkName, NodeIdx, ValueX)
      , ValueX < FunctionValue
      , NodeIdx != OldNodeIdx
    } = NewNodeIdx
  , possibleNetworkIndex(NetworkName, NewNodeIdx+Shift + 1)
  , nodeFunctionValue(NetworkName, OldNodeIdx, FunctionValue)
  , #count{OldNodeIdxB :
        nodeFunctionValue(NetworkName, OldNodeIdxB, FunctionValue)
      , OldNodeIdxB < OldNodeIdx
      , possibleNetworkIndex(NetworkName, OldNodeIdxB)
    } = Shift
  , sbn(NetworkName, _)
  , generateOrderedFunctionsVersionOfSbn(NetworkName)
.
% every and each function must be remapped
:-
    #count{ OldNodeIdx, NewNodeIdx :
         mapIndex(NetworkName, OldNodeIdx, NewNodeIdx)
    } != Dimension
  , sbn(NetworkName, Dimension)
  , generateOrderedFunctionsVersionOfSbn(NetworkName)
.
:-
    #count{ NodeIdx, FunctionValue :
          orderedNodeFunctionValue(NetworkName, NodeIdx, FunctionValue)
    } != Dimension
  , sbn(NetworkName, Dimension)
  , generateOrderedFunctionsVersionOfSbn(NetworkName)
.
% SBFs remapping
orderedSbf(NetworkName, NewNodeIdx, IneqInputs, IneqCount) :-
    sbf(NetworkName, OldNodeIdx, IneqInputs, IneqCount)
  , mapIndex(NetworkName, OldNodeIdx, NewNodeIdx)
  , generateOrderedFunctionsVersionOfSbn(NetworkName)
.
% network layout remapping
orderedNodeInput(NetworkName, NewNodeIdx, InputIdx, NewSrcNodeIdx) :-
    nodeInput(NetworkName, OldNodeIdx, InputIdx, OldSrcNodeIdx)
  , mapIndex(NetworkName, OldNodeIdx, NewNodeIdx)
  , mapIndex(NetworkName, OldSrcNodeIdx, NewSrcNodeIdx)
  , generateOrderedFunctionsVersionOfSbn(NetworkName)
.
% weight generation for every input of each node
&dom{-Dimension..Dimension} = orderedWeight(NetworkName, NodeIdx, InputIdx) :-
possibleNetworkIndex(NetworkName, NodeIdx)
, possibleNetworkIndex(NetworkName, InputIdx)
  , sbn(NetworkName, Dimension)
  , generateOrderedFunctionsVersionOfSbn(NetworkName)
.
% networks weight’s remapping
:-
    generateOrderedFunctionsVersionOfSbn(NetworkName)
  , mapIndex(NetworkName, OldNodeIdx, NewNodeIdx)
  , &sum{weight(NetworkName, OldNodeIdx, InputIdx)} = W1
  , &sum{orderedWeight(NetworkName, NewNodeIdx, InputIdx)} = W2
  , not W1 = W2
  , limitWeights(NetworkName, W1)
  , limitWeights(NetworkName, W2)
  , possibleNetworkIndex(NetworkName, InputIdx)
.
limitWeights(NetworkName, -Dimension..Dimension) :-
    sbn(NetworkName, Dimension)
.
~~~

## Acknowledgements

Sergiu Ivanov is partially supported by the Paris region via the project DIM RFSI n°2018-03 “Modèles informatiques pour la reprogrammation cellulaire”. The authors would also like to thank the IDEX program of the University Grenoble Alpes for its support through the projects COOL : this work is supported by the French National Research Agency in the framework of the Investissements d’Avenir program (ANR-15-IDEX-02). This work is also supported by the Innovation in Strategic Research program of the University Grenoble Alpes. The authors would thanks Ibrahim Cheddadi for fruitful discussions.

